# Immunolocalization of fasciclin-like arabinogalactan proteins in the G-layers of poplar tension wood fibers

**DOI:** 10.1101/2024.12.19.629400

**Authors:** Myiuki Takeuchi, Fernanda Trilstz Perassolo Guedes, Françoise Laurans, Nathalie Boizot, Annabelle Déjardin, Gilles Pilate

**Affiliations:** BioForA, INRAE ONF, 45000 Orléans, France; The Graduate School of Engineering, The University of Tokyo, Tokyo, Japan; Sylvamo Brasil, Papel e produtos florestais, Mogi-Guacu, São Paulo, Brasil

**Keywords:** Populus, antibody, arabinogalactan proteins, secondary cell wall

## Abstract

In hardwood trees, tension wood (TW) is an adaptive mechanism used by trees to orient their stems and branches, withstand their own weight, and improve wind resistance. In many species, such as poplar, TW fiber cell walls exhibit a supplemental layer, named the G-layer, which is responsible for the mechanical properties of TW. However, the molecular mechanisms involved still need to be clarified.

The synthesis of a number of fasciclin-like arabinogalactan proteins (FLA) has been shown to be highly upregulated during tension wood formation in poplar and is potentially associated with the outstanding mechanical properties of tension wood. Three polyclonal antibodies directed against different poplar TW-specific FLA epitopes were produced and used to assess the presence of these FLAs in differentiating and mature tension wood fibers.

Using immunohistochemistry, FLAs were detected at early stages of G-layer differentiation, specifically at the inner side of G-fiber cell walls, whereas a weaker signal was detected in mature G-fibers. However, western blot analyses of protein extracts from differentiating and mature tension wood revealed increased levels of FLA in mature TW fibers, suggesting that these FLAs remained present in mature G layers but were not accessible to anti-FLA antibodies in TW histological sections.

Overall, specific FLAs involved in secondary cell wall construction are located at the inner side of the G-layer and are likely actors in the unique mechanical properties of TWs.

## INTRODUCTION

Tension wood (TW) is a unique wood with outstanding mechanical properties (Alméras and Fournier 2009). It is produced by hardwood trees to orient their branches and main axes, to help support the weight of their crown and to resist prevailing winds. TW acts to maintain or restore static equilibrium among the different parts of a tree (Hartmann 1942, cited by Sinnott 1952), which is also referred to as posture control (Moulia et al. 2006). Strong maturation stresses are generated in TW via a mechanism that has not yet been elucidated (Alméras and Clair 2016). In poplar, as in many temperate hardwood species, TW consists of specific fibers (named G-fibers) that possess an extra secondary cell wall (SCW) layer named the gelatinous layer (or G-layer) deposited at the inner face of the S2 layer. This G-layer appears to be mostly made of crystalline cellulose with a very low microfibril angle. However, the G-layer also contains a matrix of noncellulosic polysaccharides containing rhamnogalacturonan I (RG-I) pectins and arabinogalactan proteins (AGPs) (Guedes et al. 2017). In addition, transcriptomic analyses performed on differentiating and mature tension wood have revealed very high and specific expression of a number of AGP transcripts in poplar TW (Déjardin et al. 2004; Lafarguette et al. 2004; Andersson-Gunnerås et al. 2006).

AGP are known to be cell surface glycoproteins with a core-protein backbone with repetitive dipeptide motifs such as Ala-Pro, Ser-Pro, Thr-Pro, and Val-Pro, where the Pro residues are converted to Hyp, which are further O-glycosylated by complex carbohydrates, usually type II arabinogalactan (AG) polysaccharides (Gaspar et al. 2001; Kieliszewski 2001; Showalter 2001). Type II AGs are made of a β-1,3-galactan backbone with β-1,6-galactosyl branches decorated with arabinosyl residues and occasionally with other sugar residues, such as glucuronic acid, rhamnose, and fucose (Tan et al. 2004). AGPs often harbor a glycosylphosphatidylinositol (GPI) lipid anchor attached to their C-terminus during posttranslational modifications, which suggests that these AGPs remain attached to the outer side of the cell membrane when the xylem fiber is still alive.

A distinct subgroup of AGPs strongly represented in TW are fasciclin-like AGPs (FLAs), which, in addition to AG modules, harbor a fasciclin-like domain (Johnson et al. 2003). Fasciclins were first described in *Drosophila* as adhesion molecules important for axonal guidance (Elkins et al. 1990). Thus, FLAs possess domains for both protein and carbohydrate interactions, which are important for their functions (Seifert 2018). Interestingly, in Arabidopsis, two FLA genes (*AtFLA11* and *AtFLA12*) have been shown to be coexpressed with *ces*A genes specific to SCW cellulose biosynthesis (Persson et al. 2005). The Arabidopsis knockout mutant for these two FLA genes presents altered stem mechanical properties, namely, reduced tensile strength, as well as altered cell wall architecture and composition, with increased cellulose microfibril angles and reduced arabinose, galactose and cellulose contents (MacMillan et al. 2010). These findings suggest that these FLAs contribute to plant stem biomechanics by affecting both cellulose deposition and the integrity of the cell wall matrix. These two glycoproteins are possible cell surface sensors that regulate SCW development in response to mechanical stresses (Ma et al. 2022).

Bioinformatic analyses of the *Populus trichocarpa* genome led to the identification of 35--50 FLA sequences according to the research criteria chosen (Zang et al., 2015; Showalter et al. 2016). Interestingly, phylogenetic analysis of the PtFLAs identified by Showalter et al. (2016) revealed that 24 of them have the highest homology with AtFLA12 and AtFLA11. In *Populus tremula × Populus alba* hybrids, 15 FLA sequences (named PopFLAs for poplar fasciclin-like arabinogalactan proteins) isolated from different xylem cDNA libraries have been shown to be expressed in the differentiating xylem once SCW formation was initiated (Lafarguette et al. 2004; Andersson Gunnerås et al. 2006). Interestingly, ten of them (PopFLA1-10) appeared to be specifically expressed in TW (Lafarguette et al. 2004) and were among the most highly represented sequences in a differentiating TW cDNA library, indicative of their high expression level (Déjardin et al. 2004). The deduced protein sequences of these 15 PopFLAs feature group A fasciclin-like AGPs, as defined by Johnson et al. (2003), with a fasciclin-like domain flanked by two AGP-like regions rich in Ala, Pro, Ser and Thr. They all present a conserved structural organization with only some variations in the length and composition of the first AGP-like domain (Lafarguette et al. 2004). Similarly, all 15 PopFLA sequences were predicted to have an N-terminal signal peptide for secretion, while thirteen of them were also predicted to have a C-terminal signal for the addition of a GPI anchor (Lafarguette et al. 2004).

In this work, we further investigated the localization of PopFLAs highly and specifically expressed during the building of G-fiber SCWs. To this end, we produced specific antibodies directed against the FLA peptide moiety. Using both immunocytochemistry and western blotting, we determined their precise localization during TW differentiation.

## MATERIAL AND METHODS

### Bioinformatic analyses

Sequence alignments were produced via ClustalW (CLUSTAL 2.1 Multiple Sequence Alignments; https://www.genome.jp/tools-bin/clustalw; Thompson et al. 1994). The occurrence of a signal peptide was predicted with TargetP-2.0 (https://services.healthtech.dtu.dk/services/TargetP-2.0/; Almagro Armenteros et al. 2019). The fasciclin-like domain (IPR00782 domain) was predicted via InterproScan (http://www.ebi.ac.uk/interpro/scan.html; Jones et al. 2014). The position of the GPI anchor signal sequence was predicted via NetGPI-1.1 (https://services.healthtech.dtu.dk/services/NetGPI-1.1/; Gislason et al. 2021). N-glycosylation sites (Asp-X-Ser/Thr) were predicted with the NetNGlyc tool v1.0 (https://services.healthtech.dtu.dk/services/NetNGlyc-1.0/; Hamby et al. 2008).

### Evolutionary analysis via the maximum likelihood method

The evolutionary history was inferred via the maximum likelihood method and the JTT matrix-based model (Jones et al. 1992). Initial tree(s) for the heuristic search were obtained automatically by applying neighbor-joining and BioNJ algorithms to a matrix of pairwise distances estimated via the JTT model and then selecting the topology with the superior log likelihood value. Evolutionary analyses were conducted in MEGA11 (Tamura et al. 2021).

### Production and characterization of anti-PopFLA antibodies

Rabbit polyclonal antibodies were generated against two FLAs specific to TW: PopFLA1 (accession # AY607753 for the full cDNA sequence in GenBank, PtXaTreH.19G106400.1.p gene model in *P. tremula × P. alba* HAP1 v5.1 genome, PtXaAlbH.19G100600.1.p gene model in *P. tremula × P. alba* HAP2 v5.1 genome, Potri.019G121200.1.p in *P. trichocarpa* v4.1 genome) and PopFLA8 (accession # AY607760, PtXaTreH.09G008100.1.p in *P. tremula × P. alba* HAP1 v5.1, PtXaAlbH.09G008100.1. in *P. tremula × P. alba* HAP2 v5.1, Potri.009G012200.1.p in *P. trichocarpa* v4.1) (http://phytozome.jgi.doe.gov/; Zhou et al. 2023). Three different antibodies were generated: anti-PopFLA1 and anti-PopFLA8 were produced using the full recombinant proteins as antigens. The third antibody (anti-PF1) was directed against a 15-amino-acid peptide (DGDKSELVKFHVVPT) from the PopFLA1 protein (Fig. 1).

**Fig. 1.**
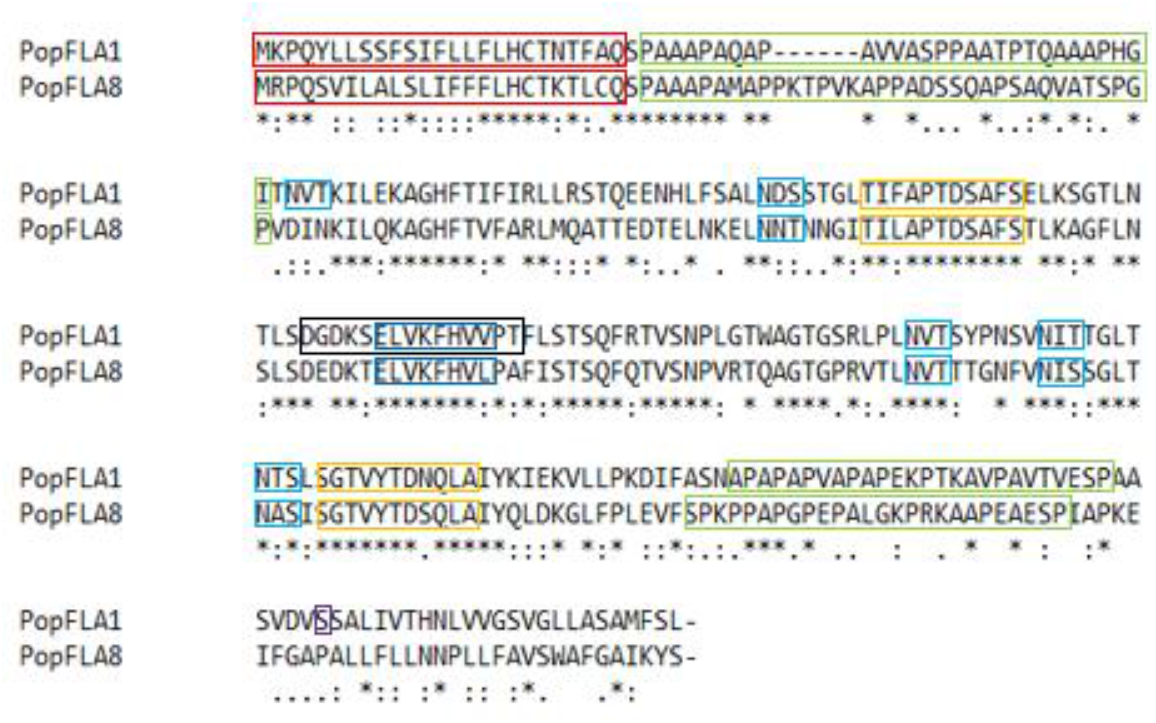
Sequence alignment of PopFLA1 and PopFLA8. Red frames: signal peptide for secretion; green frames: AGP-like domains; orange frames: the HI and H2 conserved regions characteristic of FAS1 domain; dark blue frames: amino acids involved in adhesion; light blue frames: N-glycosylation sites (Asp-X-Ser/Thr); purple frame: position of the PopFLA1 GPI anchor signal sequence; black frames: 15 amino acid peptide sequence used as an immunogen to produce the anti-PF1 antibody.

#### Production of anti-PopFLA1 and anti-PopFLA8 antibodies

To produce the PopFLA1 and PopFLA8 recombinant proteins, we choose a full-length cDNA clone for each of them: the physical clone PtaXM0028A2 (accession # CF229653 in GenBank dbEST at NCBI) for PopFLA1, with a nucleotide sequence of 1040 bp corresponding to a protein of 263 aa; the physical clone PtaXM0020E10 (accession # CF229042) for PopFLA8, with a nucleotide sequence of 1132 bp corresponding to a protein of 269 aa (for details on the acquisition of these clones, see Déjardin et al. 2004)). The whole coding sequence of each clone was PCR-amplified via the 556F and 556R primers for PopFLA1 and the 560F and 560R primers for PopFLA8 (Sup Table 1). PCR-amplified fragments were cloned and inserted into the pENTR vector via the TopoCloning Kit (Invitrogen) and transferred via LR recombination into the Gateway pDEST-17 (Invitrogen) expression vector, generating pDEST17-556 and pDEST17-560. His-tagged recombinant proteins were produced in the *E. coli* strain Rosetta pLysS (Invitrogen) supplemented with 0.1 mM IPTG. His-tagged recombinant proteins were purified in batches via immobilized metal affinity chromatography resin charged with cobalt (TALON^®^). Purified recombinant proteins were injected into 2 rabbits for the production of polyclonal antibodies (AGRO-BIO, La Ferté Saint Aubin, France): immune sera were obtained after 77 days, and five intradermal injections were given at J0, J14, J28, J42 and J56. Anti-PopFLA1 and anti-PopFLA8 antibodies were purified against their respective His-tagged recombinant proteins via activated immunoaffinity support Affi-Gel® 10 Gel (Bio-Rad).

#### Production of the anti-PF1 antibody

The 15-amino-acid peptide (DGDKSELVKFHVVPT) was synthesized by AGRO-BIO (La Ferté Saint Aubin, France) and injected into two rabbits for polyclonal antibody production. The anti-PF1 antibody was purified against the peptide via an activated immunoaffinity support Affi-Gel® 10 Gel (Bio-Rad).

#### Characterization of polyclonal antibodies

The cross-reactivity of the different antibodies against the PopFLA1 and PopFLA8 recombinant proteins was evaluated via dot-blotting. A total of 0.2 to 25 ng of recombinant protein was spotted on an Amersham Hybond-C Extra nitrocellulose membrane and then incubated first with blocking buffer containing 3% skim milk in TBS and second with purified anti-PopFLA1, anti-PopFLA8 or anti-PF1 antibodies diluted 1:500 with 3% skim milk in TBS. After washing, the sheets were incubated with monoclonal anti-rabbit IgG (γ-chain specific)-alkaline phosphatase antibody following the manufacturer’s instructions (SIGMA). The alkaline phosphatase activity was detected via BCIP/NBT (5-bromo-4-chloro-3-indolyl-phosphate/nitro blue tetrazolium) from Promega as a substrate.

### Plant material and sampling

Micropropagated poplar plants *(P. tremula × P. alba*, section Populus, clone INRA 717-1-B4) were transferred to a glasshouse, potted in 3 L pots filled with soil and individually supplied with water and fertilizers via a drip system. To induce tension wood formation, the plant stems were artificially tilted by inclining the pots at 30° from the vertical direction and maintaining the plants at this inclination angle via a rigid stick. The height and stem diameter were recorded regularly, and samples were collected three months after tilting. Samples from the differentiating (DX) and mature (MX) xylem were collected at both the upper and lower sides of the stem, which correspond to the TW and opposite wood (OW) areas, respectively. Tension wood was easily recognizable as a bright whitish crescent that made it easy to split the TW and OW samples. DX samples were lightly scraped from the debarked stem, whereas MX samples corresponded to ground wood chips collected after removal of the differentiating xylem. Wood samples were immediately frozen in liquid nitrogen and stored at −80°C before use.

### Protein extraction, purification and western blot analysis

To evaluate the presence of PopFLA1 and PopFLA8 proteins in different poplar xylem samples, total proteins were extracted via the TCA–acetone method (Damerval et al. 1986) and quantified via the Bradford method (Bradford 1976).

#### Fractionation of soluble (cytosolic) and cell wall proteins

Proteins were extracted from 1–3 g frozen samples reduced to a fine powder in 7–15 ml of buffer A (100 mM Tris-Cl pH 7.4, 2% PEG 6000, 0.2 M sodium ascorbate, 1 mM PMSF and 2% PVPP) with shaking for 1 h at 4°C. The crude extract was clarified by centrifugation at 13000 × g for 10 min at 4°C. The supernatant contained soluble cytosolic proteins (fraction A), whereas the pellet contained mostly plasma membrane-bound and cell wall-bound proteins. The pellet was washed in buffer A, centrifuged again, and re-extracted in buffer B (buffer A + 1 M NaCl) at 4°C overnight with stirring. After centrifugation, the cell wall proteins were added to the supernatant. The cell wall protein fractions were ultracentrifuged at 235000 × g for one hour. The supernatant contained the proteins weakly bound to the cell wall, whereas the membrane-bound proteins remained in the pellet. Proteins were quantified according to the Bradford method (Bradford 1976), aliquoted and stored at −20°C until use.

#### Western blot analysis

A total of 0.5 µg of protein was concentrated via the trichloroacetic acid (TCA) precipitation method. After washing, the TCA pellets were resuspended in 25 µL of 1 M Tris-HCl (pH 6.8) and 4x Laemmli sample buffer (Bio-Rad) and then separated on a 10% Mini-PROTEAN®TGX^TM^ precast gel (Bio-Rad).

Proteins were subsequently transferred onto nitrocellulose membranes via the iBlot Dry Blotting System (Invitrogen). Western blot analysis was performed using purified primary antibodies diluted at 1:10000 (anti-PopFLA1 and anti-PopFLA8 antibodies) and 1:5000 (anti-PF1 antibody) and the secondary antibody anti-rabbit IgG HRP conjugate (SIGMA) at a 1:50000 dilution. Labeling was visualized via enhanced chemiluminescence (ECL) following the manufacturer’s instructions (Amersham).

### Anatomical and histochemical analyses

#### Sample preparation

Samples were collected from the basal part of the stem of 3-month-old greenhouse-grown poplar plants. A 3 cm long stem piece was collected at the base of the tilted stem at approximately 10 cm above the ground. Wood subsamples, several mm in size (approximately 3 mm^3^), were cut from the TW and OW sides and immediately fixed for 4 h in 2.5% formaldehyde and 0.1% glutaraldehyde in 0.1 M McIlvaine citrate–phosphate buffer, pH 6.8. After dehydration in a graded series of ethanol, the samples were embedded in medium-grade LR White resin (Agar Scientific Ltd., Stansted, UK).

#### Immunolocalization by light microscopy

Semithin cross sections (0.8 µm thick) were cut via a diamond knife (Diatome, Biel, Switzerland) installed on an Ultracut R microtome (Leica, Rueil-Malmaison, France). The sections were laid on silanized slides (Dako Cytomation, Trappes, France) and fixed by heating at 54°C.

All incubations for immunolabeling were performed in a damp chamber to prevent liquid evaporation. The sections were first preincubated for 1 h with a droplet of blocking solution containing 3% bovine serum albumin (BSA) and 0.01% Tween-20 in 10 mM Tris-buffered saline (TBS), pH 7.5, and then for 10 min with droplets of dilution buffer (0.3% BSA and 0.01% Tween-20 in 10 mM TBS, pH 7.5) at room temperature. The sections were then incubated overnight at 4°C with the primary antibodies anti-PopFLA1, anti-PopFLa8 and anti-PF1 diluted 1:100 in the same buffer. The sections were then washed extensively with TBS buffer and incubated for 1 h at room temperature with 10 nm gold-conjugated secondary antibody in TBS buffer (1:50) (BB International, British-Biocell, Cardiff, UK). After 1 h, the sections were washed four times with TBS and twice with distilled water. Immunogold labeling was enhanced by incubation for 6 min with a Silver Enhancing Kit (BB International, British-Biocell, Cardiff, UK) according to the manufacturer’s instructions. The sections were then poststained with a mixture of methylene blue/Azur II (Richardson et al. 1960) and examined under a Leica DMR light microscope. To assess the specificity of labeling, controls were generated by omitting the primary antibodies.

#### Immunolocalization by electron microscopy

Ultrathin (100 nm) transverse sections of samples embedded in LR White resin were cut with an Ultracut R microtome (Leica) fitted with a diamond knife. The sections were harvested on 400 uncoated nickel mesh grids for immunocytochemical localization. The grids were allowed to float for 1 h on a blocking solution containing 3% BSA and 0.01% Tween 20 in 50 mM TBS, pH 7.5. The samples were then incubated twice for 5 min each in dilution buffer containing 0.3% BSA and 0.01% Tween 20 in 50 mM TBS, pH 7.5. The primary antibody reaction was carried out overnight at 4°C by floating the grids on drops of the primary polyclonal antibodies diluted 1:10 in the same dilution buffer. The grids were then washed extensively in TBS and incubated with 20 nm gold-conjugated secondary antibody in dilution buffer (1:50) (BB International, British-Biocell, Cardiff, UK). The grids were subsequently allowed to float for 20 min on 3% aqueous uranyl acetate solution for additional contrast and observed with a transmission electron microscope (Philips CM10) operating at 80 kV.

In each immunolabeling experiment, control sections were subjected to the treatment described above but without the addition of the primary antibody to evaluate the nonspecific binding of the secondary antibody.

## RESULTS

### Strategy for antibody production

Supplemental Figure 1 presents a phylogenetic tree built with the 15 PopFLAs identified in *P. tremula × P. alba* wood samples (Lafarguette et al. 2004) and the 28 PtFLAs from *P. trichocarpa* with the highest homology with AtFLA11 and AtFLA12 (Showalter et al. 2016). Each PopFLA is appaired to a PtFLA. PopFLA1 to PopFLA6 with homologous PtFLAs are clustered into a first group, which is well separated from a second small group that includes PopFLA8 and PopFLA9, whereas the other PopFLAs are further away in the tree. Interestingly, AtFLA11 and AtFLA12 appear to be more homologous to PopFLAs that are not specific to TW (e.g., PopFLA15, PopFLA13 and PopFLA11; Lafarguette et al. 2004).

At the amino acid level, the PopFLA9 sequence is 80% identical to PopFLA8, whereas the 13 other PopFLA sequences are at most 58% identical to PopFLA8 (Sup Table 2). PopFLA1 is part of a group of 6 very homologous genes (PopFLA1 to PopFLA6) apart from their N- and C-terminal ends (Sup Fig. 2; Lafarguette et al. 2004), with more than 82% identity when the other 9 PopFLA sequences are at most 58% identical to PopFLA1 (Sup Table 2). Interestingly, the PopFLA1 sequence appeared more homologous with the PopFLA15 sequence, which is expressed in NW and OW as well as in TW, with 58% identity, than with a number of TW-specific PopFLAs (namely, PopFLA7 to PopFLA10, with 44 to 54% identity). Likewise, in addition to PopFLA9, PopFLA15 presented the highest identity with PopFLA8. Notably, PopFLA12 presented the lowest similarity with all the other PopFLAs, with values of less than 36%.

With respect to this comparison between sequences, we chose to produce polyclonal antibodies directed against two TW-specific PopFLAs: PopFLA1 and PopFLA8. Despite their overall similarity, these two PopFLAs present some important differences (Fig. 1).

PopFLA1 is orthologous to Potri.019G121200 in the *P. trichocarpa* genome sequence and to the PtFLA29 protein sequence according to the classification of Showalter et al. (2016). Similarly, PopFLA8 is orthologous to Potri.009G012200 in the *P. trichocarpa* genome sequence and to the PtFLA2 protein sequence (Sup Fig. 1). Their protein domains are arranged as in the Arabidopsis PopFLAs group A, with a single fasciclin domain flanked by two AGP glycomodules (Johnson et al. 2003, Fig. 1). Both proteins harbor a signal peptide for secretion at their N-terminal end and two AGP-like domains with repetitive dipeptide motifs, such as Ala-Pro, Ser-Pro, Thr-Pro, and Val-Pro, where the Pro residues are further converted to Hyp to be arabinogalactosylated (Kieliszewski and Shpack 2001). The first AGP module is slightly shorter in the PopFLA1 sequence. Both proteins possess HI and H2 conserved regions characteristic of the FAS1 domain (Kawamoto et al., 1998) as well as an amino acid sequence (ELVKFHVV/L) involved in adhesion to integrin-like proteins (Tyr-His or Phe-His with Ile, Leu and Val as neighbors; Kim et al. 2000; Kim et al. 2002, Fig. 1). PopFLA1 has 4 (and possibly 5) N-glycosylation sites (Asp-X-Ser/Thr), whereas PopFLA8 has only 3 (and unlikely 4) N-glycosylation sites: indeed, the probability that the NITT site is an N-glycosylation site is rather low (4/9) for PopFLA1, and the probability for the corresponding site NISS in PopFLA8 is even lower. Finally, PopFLA1 is predicted to possess a GPI modification site (P>0.7), whereas PopFLA8 (as PopFLA6) does not. Overall, the two sequences present 50% (129/263) identity, but the identity increases to 76% (90/118) in the FAS1 domain.

### Analyses of antibody specificities

We tested the reactivity of the three antibodies against the PopFLA1 and PopFLA8 recombinant proteins (Fig. 2). The anti-popFLA1 antibody detected 1 ng of PopFLA1 and 5 ng of PopFLA8. The anti-popFLA8 antibody detected 0.2 ng of PopFLA8 and 1 ng of PopFLA1. The anti-PF1 antibody was able to detect 0.2 ng of PopFLA1 and 1 ng of PopFLA8. This indicates that the three antibodies were most likely able to recognize with different efficiencies motifs present on PopFLA1 and PopFLA8 and their close homologs, which are PopFLA2 to PopFLA6 and PopFLA9 (as shown in Sup Table 2).

**Fig. 2.**
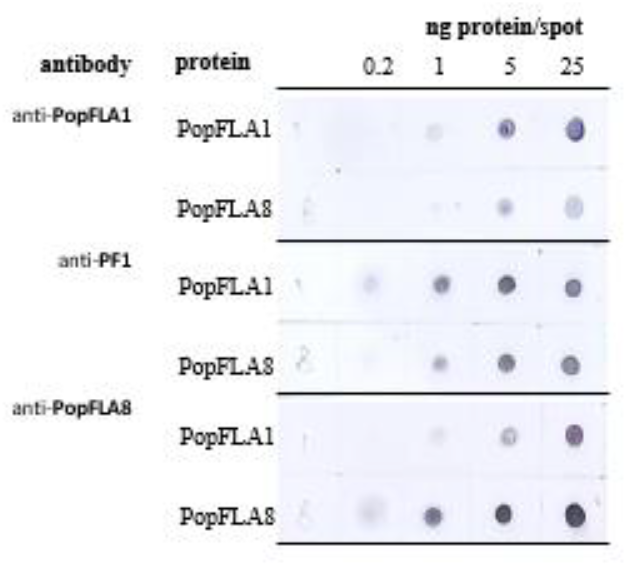
Comparison of the reactivities of three polyclonal antibodies against the PopFLA1 and PopFLA8 recombinant proteins. Two of them (anti-PopFLA1 and anti-PopFLA8) are directed against the PopFLA1 and PopFLA8 recombinant proteins, respectively; the anti-PF1 antibody is directed against a 15-amino acid peptide from the PopFLA1 fasciclin domain that encompasses its adhesion domain to integrin-like proteins (see Fig. 1). A third polyclonal antibody was directed against the 15 amino acid PF1 peptide DGDKSELVKFHVVPT, which is part of the fasciclin domain of PopFLA1 and spans the amino acids potentially involved in the adhesion of the fasciclin domain to integrin-like proteins (in bold in the formula; Fig. 1; Kim et al., 2002). When the comparison of the 15 PopFLA sequences was limited to this PF peptide sequence (Sup Fig. 3), the PF1 to PF6 sequences were completely identical, while PF8 and PF9 were 73% identical to PF1, and all the other PF sequences were less than 54% identical to PF1 (Sup Table 2).

The three antibodies were further tested against total protein extracts from differentiating (DX) and mature (MX) xylem sampled at the upper face (TW side) and lower face (OW side) of stems from inclined trees (Fig. 3). All three antibodies strongly labeled TW with no or very light signals in OW. Both anti-popFLA1 and anti-popFLA8 antibodies produced much stronger and wider signals in MX than in DX. In the DX samples, anti-popFLA1 recognized a 66 kDa protein, whereas anti-popFLA8 labeled a slightly smaller band at 60 kDa. Interestingly, the anti-popFLA8 antibody also recognized a small (less than 30 kDa), very lightly (or not) glycosylated protein in the DX of both TW and OW. The anti-PF1 antibody strongly labeled both differentiating and mature TWs: it recognized glycosylated proteins with sizes of 60–66 kDa and 42–45 kDa. A fainter band of greater size (>100 kDa) was visible in the DX of both TW and OW. With this third antibody, the signal intensity was of the same magnitude in the DX-TW and MX-TW samples, whereas faint signals of the same size were observed in the DX-OW samples.

**Fig. 3.**
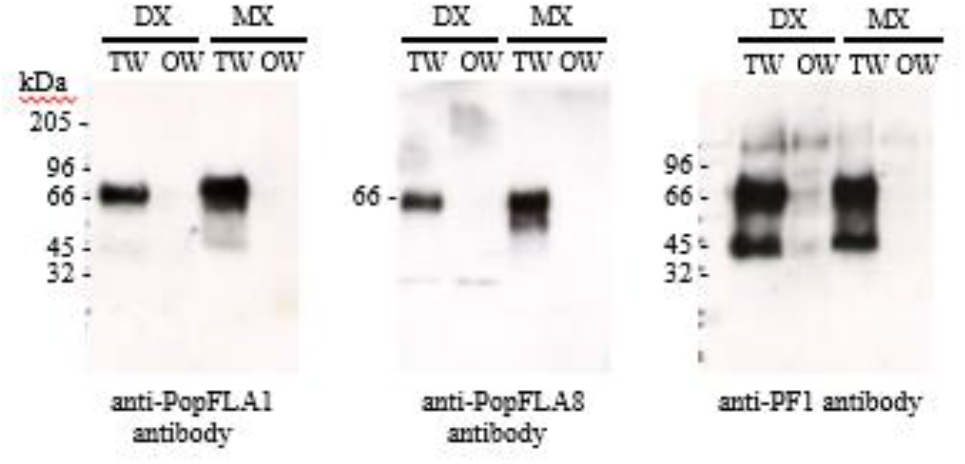
Western blot analysis of protein extracts from differentiating xylem (DX) or mature xylem (MX) of tension wood (TW) or opposite wood (OW) in a tilted poplar stem with three antibodies: anti-PopFLA1, anti-PopFLA8 and anti-PF1. Molecular mass standards (kDa) are shown on the left.

Protein extracts from differentiating and mature TWs were further fractionated into lightly bound cell wall proteins (A) and membrane-bound proteins (B). Only the membrane fraction (P) was labeled with anti-PopFLA1 and anti-PopFLA8 antibodies (Fig. 4). The anti-popFLA1 antibody detected 2 bands at 66 kDa and 40 kDa. In MX TW, the band at 40 kDa was as strong as the 66 kDa band, whereas this 40 kDa band remained faint in DX TW. Using anti-PopFLA8, we detected a strong fuzzy band at 50–60 kDa with a lighter smear at approximately 30 kDa. As with anti-PopFLA1, the signal was stronger in mature xylem than in differentiating xylem. In comparison, the JIM14 antibody, which is directed against a glycosylated AGP epitope (Knox et al. 1991), revealed a strong signal that appeared as a smear, which is typical of glycosylated proteins, in the membrane-associated fraction at high MWs (100–150 kDa, Fig. 4). In contrast to those of the other two antibodies, the signal was stronger in DX-TW than in MX-TW.

**Fig. 4.**
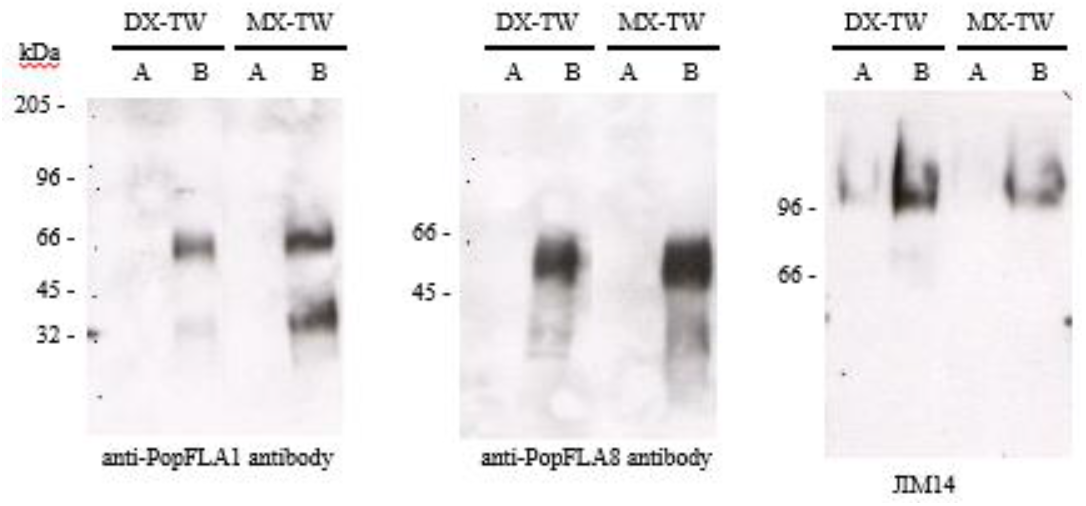
Western blot analysis of the lightly bound protein fraction (A) and the membrane-rich fraction (B) of protein extracts from tension wood (TW) differentiating xylem (DX) and mature xylem (MX) with three antibodies: anti-PopFLA1, anti-PopFLA8 and JIM14. PopFLA1 and PopFLA8 in TW xylem were linked mainly to the membrane-rich fractions. Molecular–mass standards are shown on the left.

### Localization of fasciclin-like AGP in tilted poplars

The localization of the epitopes recognized by the antibodies directed against PopFLA1, PopFLA8 and the PF1 peptide was determined on cross-sections prepared from the stems of tilted poplars producing TW and OW. To this end, we used an indirect immunolocalisation technique with colloidal gold-labeled secondary antibodies. For transmission electron microscopy (TEM), 10 nm colloidal gold particles were directly visible. In contrast, for optical microscopy observations, the gold colloid label was silver-enhanced by the precipitation of metallic silver to obtain a high contrast signal visible under a light microscope.

#### Optical microscopy observations

The labeling of the 3 antibodies was observed from the cambial zone and along the gradient of xylem differentiation in the TW (Fig. 5) and OW areas. In the first G-fibers formed in response to plant tilting (area 1), close to the cambial zone, the 3 antibodies labeled the inner faces of the young, thin G layers with varying intensities: PopFLA8 presented the strongest labeling, and PopFLA1 presented the weakest labeling (Fig. 5a, d, g). However, the labeling profiles of the 3 antibodies evolved differently in areas 2 and 3, when the G layers thickened. The PopFLA1 antibody still specifically labeled the inner face of the G layers both in area 2 and 3 (Fig. 5b, c). The inner face of the G-layers in area 2 was also strongly labeled by the PopFLA8 antibody (Fig. 5 e), but this antibody labels both the inner face and, to a lesser extent, the thickness of the differentiated G layers in area 3 (Fig. 5f). Finally, the anti-PF1 antibody labeled both the inner face and the thickness of the G layer in area 2 (Fig. 5h), and the labeling at the inner face disappeared and remained at the thickness of the differentiated G layers (Fig. 5i). We did not observe any labeling in OW (data not shown).

**Fig. 5.**
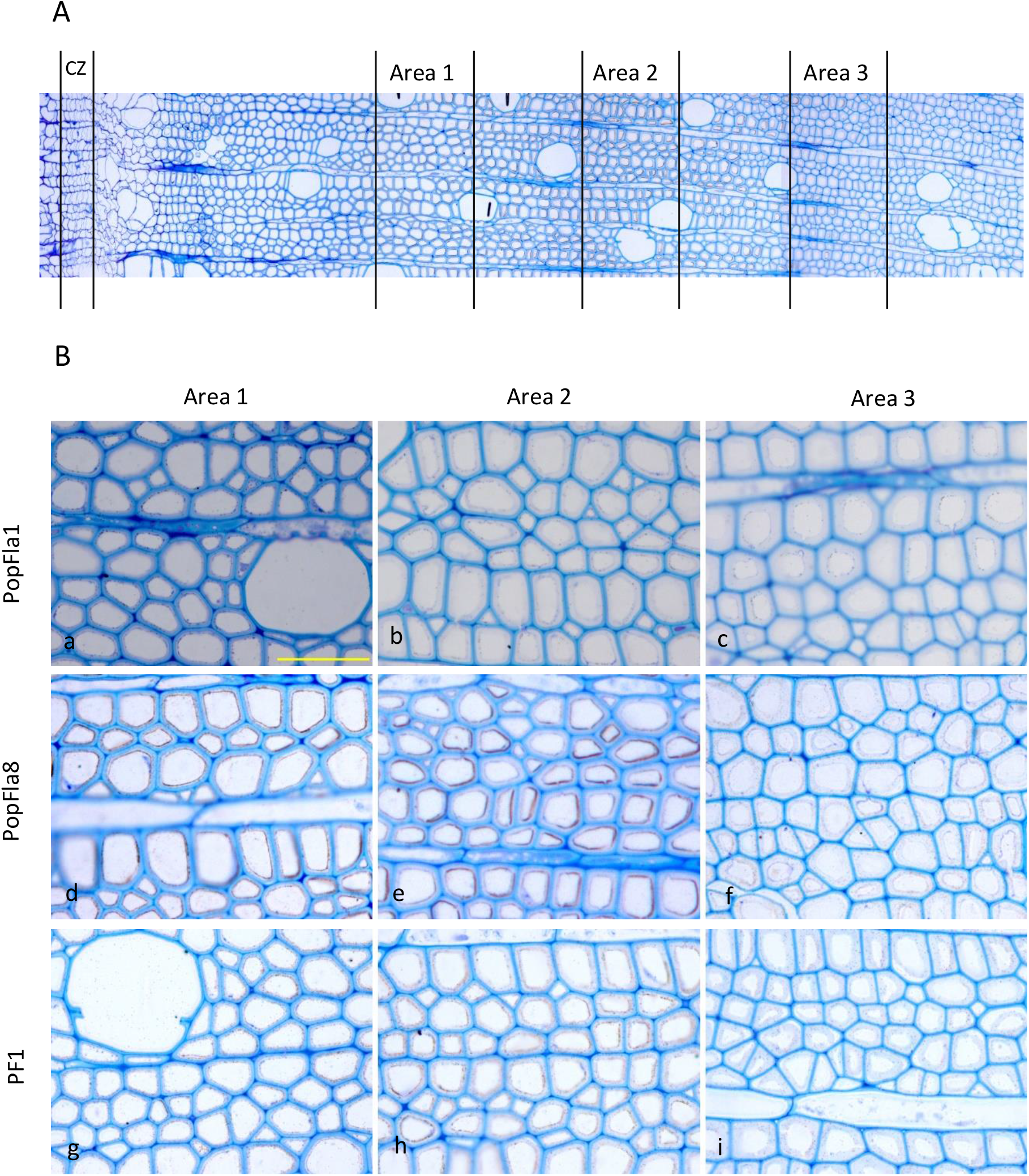
Immunogold labeling of TW with anti-popFLA1, anti-PopFLA8 and anti-PF1 antibodies observed under an optical microscope. A-Cross-sectional profiles and locations of the 3 observation areas vs the cambial zone (CZ). B-Labeling evolution along the gradient of differentiation of the G fibers. a, b, c with anti-PopFla1. d, e, f with anti-PopFla8. g, h, i with anti-PF1. Coloration Richardson’s (bar = 50 µm).

#### TEM observations

TEM observations were conducted on 2 areas with G-fibers at two different stages of differentiation: DX with thin G-layers and MX with thicker G-layers. TEM observations confirmed and refined the pattern observed via optical microscopy.

Using anti-PopFLA1 and anti-PopFLA8 antibodies, the labeling was restricted to the G-layer and was much stronger in differentiating TW fibers (Fig. 6c, e and Fig. 7c), and the gold particles appeared to be attached mainly to the plasma membrane (Fig. 6c, arrows) at the inner face of the fibers. The signal obtained with these 2 antibodies decreased during G-fiber maturation. With anti-PopFLA1, the label remained present, although weaker, at the inner face of the G-layer (Fig. 6d, f). In the case of anti-PopFla8, labeling was also present inside the G layer (Fig. 7d), while some labeling may also have occurred in the other cell wall layers. The anti-PF1 antibody labeled both the inner face of the differentiating G-layers and, to a lesser extent, the G-layer thickness (Fig. 8c, e). This labeling remained present throughout the entire thickness of the mature G layers (Fig. 8 d, f). No labeling was observed in DX-OW or MW-OW with any of the three antibodies (Figs. 6--8).

**Fig. 6.**
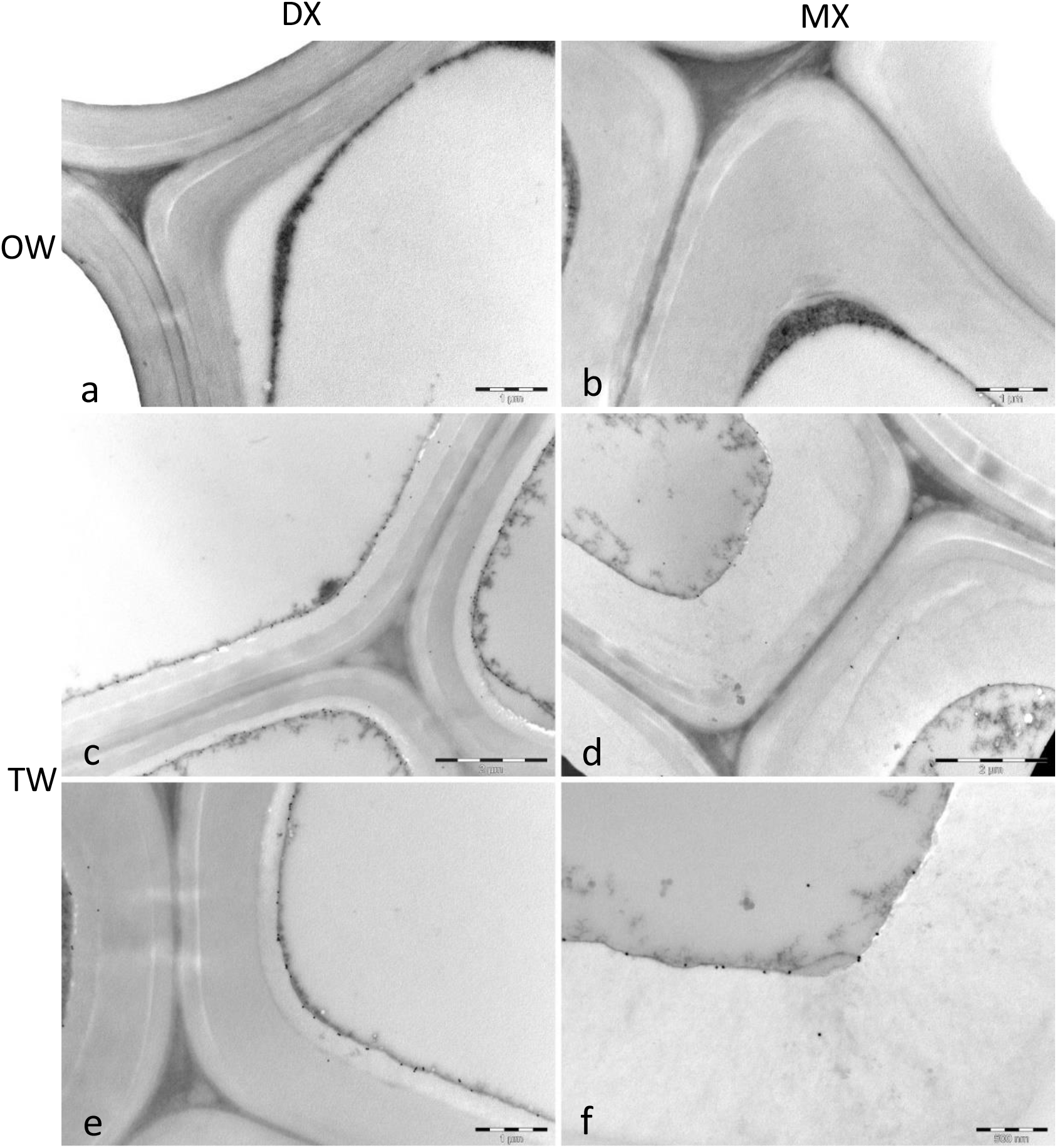
Cellular localization of anti-PopFLA1 epitopes in transverse ultrathin sections of a bent poplar stem analyzed via transmission electron microscopy. a, b Opposite wood fiber cell walls. c-f Tension wood fiber cell walls. a, c, e Differentiating G-fibers. b, d, f Mature G-fibers.

**Fig. 7.**
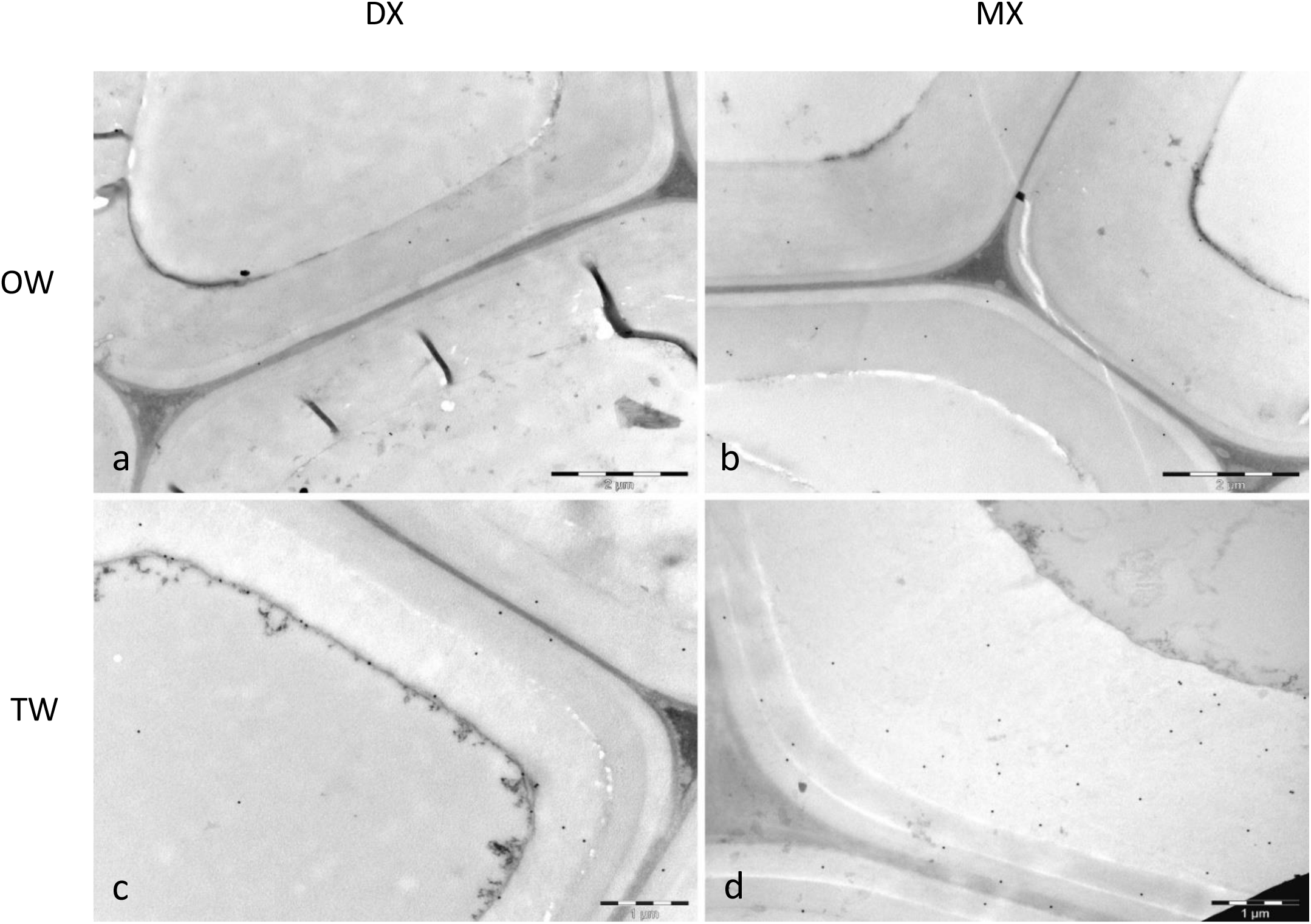
Cellular localization of anti-PopFLA8 epitopes in transverse ultrathin sections of a bent poplar stem analyzed via transmission electron microscopy. a, b Opposite wood fiber cell walls. c, d Tension wood fiber cell walls. a, c Differentiating fibers. b, d Mature fibers.

**Fig. 8.**
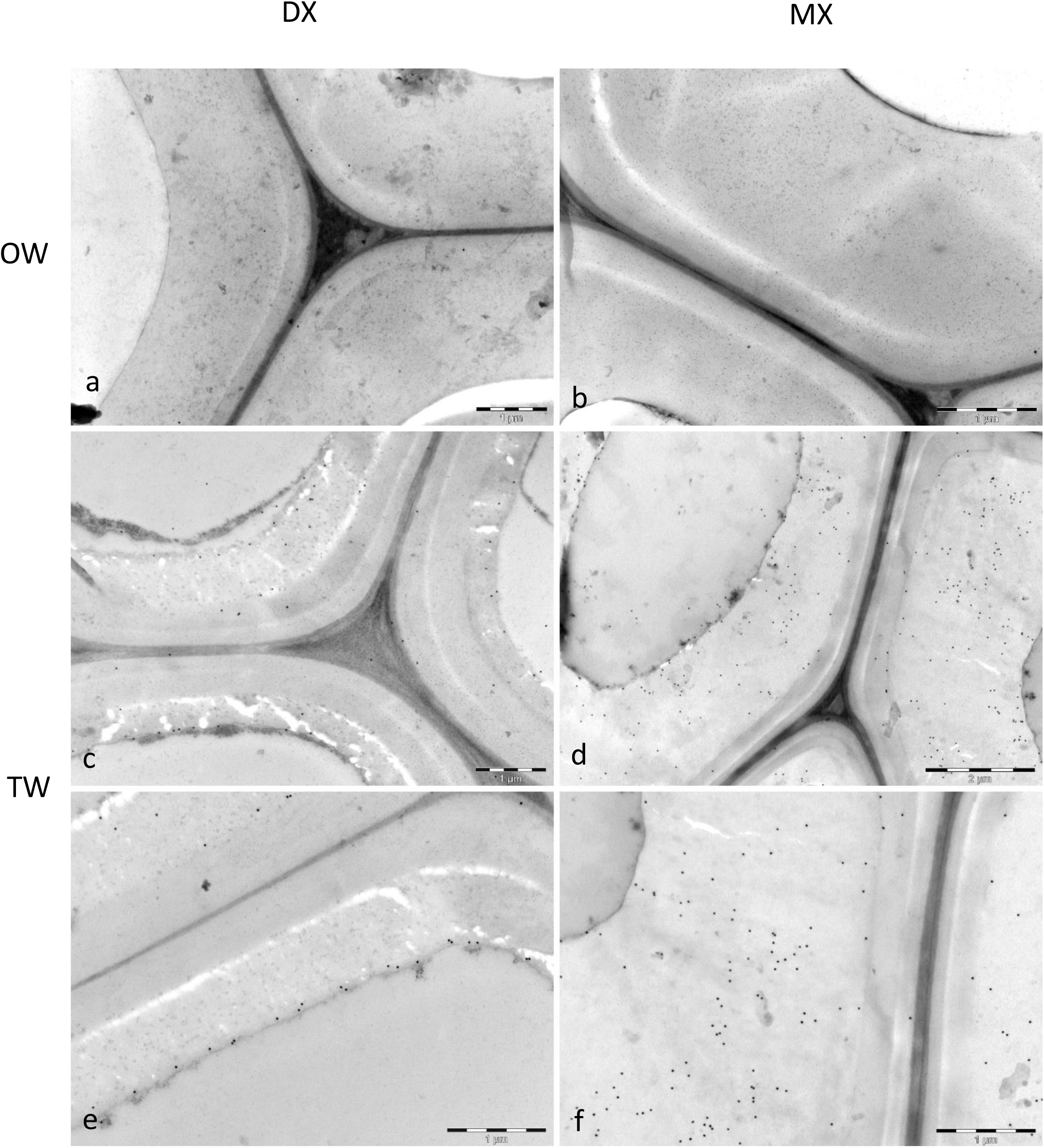
Cellular localization of anti-PF1 epitopes in transverse ultrathin sections of a bent poplar stem analyzed via transmission electron microscopy. a, b Opposite wood fiber cell walls. c-f Tension wood fiber cell walls. a, c, e Differentiating fibers. b, d, f Mature fibers.

## DISCUSSION

We have generated rabbit polyclonal antibodies against two fasciclin-like arabinogalactan proteins, PopFLA1 and PopFLA8. Genes encoding these two proteins are highly expressed in differentiating and mature xylem in tension wood, whereas they are not expressed in cambium or in OW xylem (Déjardin et al. 2004; Lafarguette et al. 2004). These two PopFLA proteins belong to a group of ten proteins specifically expressed in tension wood, whereas five others are present both in tension wood and opposite wood. We chose to produce antibodies directed against the peptide parts of these glycoproteins to be able to specifically localize them during TW formation. Our antibodies are directed against 2 recombinant proteins (anti-PopFLA1 and anti-PopFLA8) or a peptide (anti-PF1) with no glycosylation, as they were produced in bacteria. There are very few antibodies directed against the peptide part of FLA. Shi et al. (2003) produced polyclonal antibodies directed against a synthetic peptide sequence within the first fasciclin domain of Salt Overly Sensitive5, a putative cell surface adhesion protein required for normal cell expansion in Arabidopsis. Similarly, Wang et al. (2015) produced a polyclonal antibody directed against PopFLA6 (named PtFLA6 in their paper, equivalent to PtFLA39 in Showalter notation (see Showalter et al. 2016 and Sup Fig. 1, this paper)).

### These antibodies are specific to TW PopFLA

Both anti-PopFLA1 and anti-PopFLA8 antibodies recognized PopFLA in DX-TW and MX-TW, but no or very faint signals were detected in OW by western blotting or immunocytochemistry (Figs. 3-7); these findings strongly suggest that both antibodies recognize only or merely TW-specific FLA. The absence of labeling in the DX-OW and MX-OW samples (Figs. 3, 6, 7) indicates that both antibodies do not recognize epitopes on PopFLA11 to PopFLA15, which are not regulated between TW and OW (Lafarguette et al. 2004). Owing to the homologies between the sequences of the different PopFLAs expressed in poplar wood (Sup Table 2), we anticipate that they recognized PopFLA1-6 and PopFLA8-9 well (Fig. 2).

### An effect of the presence of a GPI anchor on PopFLA localization?

The presence of TW-specific PopFLA1 at the inner side of the G-layer, which is also the outer surface of the plasma membrane in living fibers, is in accordance with the sequence-based prediction of GPI anchorage for PopFLA1 (Fig. 6c). In MX TW, the labeling pattern clearly differed between PopFLA1 and PopFLA8: the anti-PopFLA1 antibody still labeled the inner side of the mature G-layer (Fig. 6), whereas the anti-PopFLA8 antibody labeled the whole thickness of the G-layer (Fig. 7). This may be related to PopFLA8 being (as PopFLA6) a PopFLA sequence with no predicted GPI anchor to the plasma membrane. Similarly, Wang et al. (2017), using their anti-PopFLA6 antibody (named PtFLA6 in their paper), observed a label sprayed on the thickness of the G layer, in accordance with the absence of a predicted GPI anchor in PopFLA6. The anti-PopFLA8 antibody lightly labeled the other layer of the cell wall, both in TW and OW; this may be due to unspecific labeling, although we cannot rule out that it recognized epitopes present on a wider range of PopFLAs.

### Previous anti-AGP antibodies were directed against the glycosidic part of the AGP

Several antibodies that recognize polysaccharide motifs specific to AGP have been produced using the purified AGP fraction as an immunogen and are still widely used today (e.g., Guedes et al. 2017). However, most of the time, the epitopes recognized by these antibodies are not always identified, and their ability to recognize members of different AGP families is not known. In fact, we do not know if any of these antibodies recognize FLA. For example, the AGP epitope of JIM14 was initially described as an unknown oligosaccharide sequence present on AGP associated with both the plasma membrane and the cell wall (Knox et al. 1991). JIM14 strongly labels high-MW proteins (110 and 200 kDa) in TW, with strong labeling in MX-TW and even stronger labeling in DX-TW, while a mild signal is detected in OW samples (Lafarguette et al. 2004; Fig. 4). In histological sections, JIM 14 labels fibers as well as vessels and rays; in fibers, the labeling is concentrated on the primary cell wall and middle lamellae, while strong labeling is also present at the inner face of the G-layer in TW fibers (Lafarguette et al. 2004). Taken together, the discrepancies between the labeling patterns of JIM14 and antiPopFLA1 and antiPopFLA8 antibodies suggest that JIM14 does not recognize FLA oligosaccharide motifs but probably recognizes motifs present on a number of TW-regulated AGPs as well as on other AGPs present in the primary cell wall and middle lamella of xylem cells.

### These antibodies are directed against the peptide part of popFLA

PopFLA1 has 263 amino acids, which theoretically results in a 27 kDa peptide, whereas PopFLA8 has 269 amino acids, which theoretically results in a 29 kDa peptide. The anti-PopFLA1 antibody recognized a 66 kDa band, the anti-PopFLA8 antibody recognized a slightly smaller band, and the anti-PF1 antibody labeled merely 40--70 kDa. These findings indicate that the PopFLAs detected in TW may not be highly glycosylated. The size of the proteins recognized by the three antibodies is far smaller than that of the AGP detected by antibodies directed against the glycosidic part of AGP, such as JIM14 (110 kDa and 200 kDa) in poplar TW samples (Lafarguette et al. 2004; Fig. 4). However, we cannot rule out that more highly glycosylated PopFLA, if present, may not be recognized by our antibodies.

### Two major bands of different sizes and intensities are consistently detected by either of the 3 antibodies

The third antibody is directed against the PopFLA1 fasciclin domain, more precisely, a 15 amino acid sequence, named PF1, which spans the amino acids potentially involved in the adhesion of the fasciclin domain to integrin-like proteins, as shown in another model (Kim et al. 2002). This sequence is very well conserved in a number of TW-specific PopFLA sequences (PopFLA1 to 6 with 100% identity and PopFLA8 & PopFLA9 with 73% identity) and at most 53% identity for FLA not specific to TW (PopFLA15 & PopFLA14). The epitopes labeled by the anti-PF1 antibody appeared rather specific to tension wood, even though some weak labeling was detected in DX-OW (Fig. 3). Similarly, a third band of large size (>100 kDa) but rather faint compared with the other bands appeared equally labeled in DX-TW and DX-OW and was barely visible in MX-OW; therefore, this antibody recognizes, to some extent, some PopFLA known to be expressed in OW, such as PopFLA15, which shares rather high homology with PopFLA1, PopFLA8 and PF1 (Sup Table 2). Interestingly, in addition to the strong band detected at 60--70 kDa, which is approximately the same size that is recognized by anti-PopFLA1 and anti-PopFLA8 antibodies, the anti-PF1 antibody also recognized a band below 45 kDa in both DX-TW and MX-TW. We may wonder whether these 2 strong bands detected by the anti-PF1 antibody may be linked to the binding of PopFLA to other proteins, such as integrins, as shown in other studies (Kim et al. 2002), or to other PopFLAs through their fasciclin domain: in this case, the 40 kDa band may correspond to PopFLA alone when the 70 kDa band results from the attachment of PopFLA to another protein. It would be interesting to determine if these two bands are those detected by anti-FLA1 and anti-FLA8 antibodies in the membrane-rich fraction of MX-TW (Fig. 4). The largest band at 60--70 kDa is consistently highly labeled by either antibody, whereas the labeling of the smaller band (below 45 kDa) is highly variable from very light to very strong intensity. We anticipate that the variations in the relative intensities of these two bands reflect the associations of PopFLA with other proteins during cell differentiation and cell wall formation.

### TW-PopFLA in TW-MX vs TW-DX: contrasting results between western blot and immunohistology

Western blots with either anti-PopFLA1 or anti-PopFLA8 antibodies revealed stronger and thicker labeling in MX-TW than in DX-TW (Fig. 3), possibly because of the greater amount of PopFLA and wider diversity in the glycosylation pattern of PopFLA in mature TW fibers. This was also observed at the RNA level: Lafarguette et al. (2004) reported that the expression level of PopFLA11, 13, 14 and 15 transcripts was greater in differentiating xylem than in mature xylem both in TW and OW, whereas the reverse was observed for PopFLA1-10 but only in TW. A similar evolution was observed by Andersson-Gunnerås et al. (2006) in their own study. In contrast, there was not much difference in the intensity of the bands labeled with the anti-PF1 peptide between DX-TW and MX-TW (Fig. 3). Immunolocalization studies indicate that TW-specific PopFLAs are bound to the plasma membrane and the cell wall. Anti-PopFLA1 and anti-PopFLA8 antibodies produced similar results: strong labeling only or merely at the inner side of differentiating G-layers with lighter labeling in mature G-layers (Fig. 5, Fig. 6 and Fig. 7). This finding clearly contradicts the results obtained by western blot. We hypothesize that PopFLA may be embedded in mature xylem by other cell wall compounds such that the anti-PopFLA antibodies can no longer reach their epitopes. The nature of these cell wall compounds remains to be determined.

One possibility is that mature PopFLAs aggregate via their fasciclin domain or interact with other cell wall proteins to strengthen the cell wall, as has been proposed for SOS5 (Shi et al. 2003, Griffiths et al. 2014).

In addition, PopFLAs may interact via their glycosylated AGP modules with some polysaccharides, such as RG-I pectins, which are very abundant in the G layer, as previously reported (Gorshkova et al. 2015; Guedes et al. 2017). Keegstra et al., in 1973, proposed that rhamnose residues on type II AG side chains may be attached to RG I. More recently, an AGP has been shown to be covalently linked to rhamnosyl residues of RG I and with arabinoxylan polysaccharides to form a network that contributes to cell wall strength (Tan et al. 2013). The existence of a protein–glycan network somewhat analogous to the peptido–glycan network of bacterial cell walls (as proposed by Lamport et al. in 1970) is in accordance with our observations: indeed, the reduction/disappearance of the immunolabeling signal in mature TW fibers (Figs. 5, 6, 7) may result from the appearance of this network, which may prevent the recognition of epitopes by the antibodies, whereas this signal remains very strong on western blots of the MX TW protein extracts (Figs. 3, 4).

Therefore, TW-specific PopFLAs may act as structural proteins involved in a supramolecular network in association with other cell wall components that may contribute to the integrity of cell walls, as proposed by Hijazi et al. (2014) for AGP31 in Arabidopsis. Alternatively, the appearance of this network in mature TW fibers may induce the formation of a gel, leading to the generation of tensile maturation stress in G-layer cellulose microfibrils, which is important for the development of the peculiar mechanical properties of TW (Alméras and Clair 2016).

## Conclusion

Anti-PopFLA1, anti-PopFLA8 and anti-PF1 peptide antibodies exhibited good specificity toward TW-specific PopFLA. These PopFLA proteins appear to be less heavily glycosylated than previously thought. PopFLA may be bound to other components of the G-layer during its differentiation and form a network. Overall, these three antibodies will be very useful for investigating the still elusive functions of these proteins during tension wood formation.

## Acknowledgments

Some of the histology observations were performed at the Tree Cell Engineering Laboratory (LICA, https://www6.inrae.fr/in-sylva-france_eng/Services/In-Lab/LICA). Miyuki Takeuchi was funded by a Haigneré fellowship (INRA). Fernanda Guedes was funded by CAPES (Ministry of Education, Brazil). We thank GBFOR (INRAE, Forest Genetics and Biomass Facility), https://doi.org/10.15454/1.5572308287502317E12, for managing the plant material in the greenhouse.

**Sup Table 1** Oligonucleotide sequences used to construct the expression vectors for recombinant protein production.

**Sup Table 2** Identity at the amino acid level between the fifteen PopFLAs expressed in the xylem and the three sequences used as immunogens to produce each of the 3 antibodies directed against the PopFLA1, PopFLA8 and PF1 peptides. For each PopFLA, the gene models for both the *Populus tremula* and the *Populus alba* alleles are indicated according to the genome sequences available at https://phytozome-next.jgi.doe.gov/. In bold, the haplotype(s) that are 100% homologous to the transcripts present in xylem tissues; red: sequences with 73 to 100% identity between the PopFLA sequence and the immunogen sequence; green: 41 to 58% identity.

**Sup Fig. 1** Phylogenetic tree built via the tree joining method in MEGA. The tree with the highest log likelihood (−10872,55) is shown. The data are drawn to scale, with branch lengths measured in terms of the number of substitutions per site. This analysis involved 45 amino acid sequences: AtFLA11 and AtFLA12 sequences, 15 PopFLAs identified in *P. tremula × P. alba* wood samples (Lafarguette et al., 2004) and the 28 PtFLAs from *P. trichocarpa* with the highest homology with AtFLA11 and AtFLA12 (Showalter et al., 2016). There was a total of 301 positions in the final data set. Blue frame: PopFLA1 and PopFLA8 used as immunogens for antibody production; red frame: the 13 remaining PopFLAs; green frame AtFLA11 and AtFLA12.

**Sup Fig. 2** Sequence alignment of PopFLA1 to PopFLA6. Red frame: signal peptide for secretion; green frames: AGP-like domains; yellow frames: HI and H2 conserved regions characteristic of the FAS1 domain; dark blue frame: amino acids involved in adhesion; light blue frames: N-glycosylation sites; purple frame: positions of GPI anchor signal sequences not present in PopFLA6.

**Sup Fig. 3** Multiple alignment of the PF peptide (15 aa) among the 15 PopFLAs found in poplar xylem (a) and the 8 most homologous TWs specific to PopFLA (b). * conserved amino acid among all the aligned sequences; : conservative substitution.

## Reference list

Almagro Armenteros JJ, Salvatore M, Emanuelsson O, Winther O, von Heijne G, Elofsson A, Nielsen H (2019) Detecting sequence signals in targeting peptides using deep learning. Life Sci Alliance 2:e201900429. 10.26508/lsa.201900429

Alméras T, Clair B (2016) Critical review on the mechanisms of maturation stress generation in trees. J. R. Soc. Interface 13: 20160550. 10.1098/rsif.2016.0550

Alméras T, Fournier M (2009) Biomechanical design and long-term stability of trees: morphological and wood traits involved in the balance between weight increase and the gravitropic reaction. J Theor Biol 256:370–381. 10.1016/j.jtbi.2008.10.011

Andersson-Gunnerås S, Mellerowicz EJ, Love J, Segerman B, Ohmiya Y, Coutinho PM, Nilsson P, Henrissat B, Moritz T, Sundberg B (2006) Biosynthesis of cellulose-enriched tension wood in Populus: global analysis of transcripts and metabolites identifies biochemical and developmental regulators in secondary wall biosynthesis. Plant J 45:144–165. 10.1111/j.1365-313X.2005.02584.x.

Bradford MM (1976) A rapid and sensitive method for the quantitation of microgram quantities of protein utilizing the principle of protein-dye binding. Anal Biochem 72:248–254. 10.1006/abio.1976.9999

Damerval C, de Vienne D, Zivy M, Thiellement H (1986) Technical improvements in two-dimensional electrophoresis increase the level of genetic variation detected in wheat seedling proteins. Electrophoresis 7:52–54.

Déjardin A, Leplé J-C, Lesage-Descauses M-C, Costa G, Pilate G (2004) Expressed sequence tags from poplar wood tissues - a comparative analysis from multiple libraries, Plant Biol 6:55–64. 10.1055/s-2003-44744

Elkins T, Zinn K, McAllister L, Hoffmann FM, Goodman CS (1990) Genetic analysis of a Drosophila neural cell adhesion molecule: interaction of fasciclin I and Abelson tyrosine kinase mutations. Cell 60:565–75. 10.1016/0092-8674(90)90660-7

Gaspar Y, Johnson KL, McKenna JA, Bacic A, Schultz CJ (2001) The complex structures of arabinogalactan-proteins and the journey towards understanding function. Plant Mol Biol 47:161–76.

Gíslason MH, Nielsen H, Almagro Armenteros JJ, Rosenberg Johansen A (2021) Prediction of GPI-anchored proteins with pointer neural networks. Curr Res Biotechnol 3:6–13. 10.1016/j.crbiot.2021.01.001

Gorshkova T, Mokshina NE, Chernova TE, Ibragimova NN, Salnikov VV, Mikshina PV, Tryfona T, Banasiak A, Immerzeel P, Dupree P, Mellerowicz E (2015) Aspen tension wood fibers contain β-1,4-galactans and acidic arabinogalactans retained by cellulose microfibrils in gelatinous walls. Plant Physiol 169:2048–2063. 10.1104/pp.15.00690

Griffiths JS, Tsai AY, Xue H, Voiniciuc C, Sola K, Seifert GJ, Mansfield SD, Haughn GW (2014) SALT-OVERLY SENSITIVE5 mediates Arabidopsis seed coat mucilage adherence and organization through pectins. Plant Physiol 165:991–1004. 10.1104/pp.114.239400

Guedes FTP, Laurans F, Quemener B, Assor C, Laine-Prade V, Boizot N, Vigouroux J, Lesage-Descauses MC, Leplé JC, Déjardin A, Pilate G, (2017) Non-cellulosic polysaccharide distribution during G-layer formation in poplar tension wood fibers: abundance of rhamnogalacturonan I and arabinogalactan proteins but no evidence of xyloglucan. Planta 246:857–878. 10.1007/s00425-017-2737-1

Hamby SE, Hirst JD (2008) Prediction of glycosylation sites using random forests. BMC Bioinformatics 9:500. 10.1186/1471-2105-9-500

Hijazi M, Roujol D, Nguyen-Kim H, Del Rocio Cisneros Castillo L, Saland E, Jamet E, Albenne C (2014) Arabinogalactan protein 31 (AGP31), a putative network-forming protein in Arabidopsis thaliana cell walls? Ann Bot 114:1087–97. 10.1093/aob/mcu038

Johnson KL, Jones BJ, Bacic A, Schultz CJ (2003) The fasciclin-like arabinogalactan proteins of Arabidopsis. A multigene family of putative cell adhesion molecules. Plant Physiol 133:1911–1925. 10.1104/pp.103.031237

Jones P, Binns D, Chang HY, Fraser M, Li W, McAnulla C, McWilliam H, Maslen J, Mitchell A, Nuka G, Pesseat S, Quinn AF, Sangrador-Vegas A, Scheremetjew M, Yong SY, Lopez R, Hunter S (2014) InterProScan 5: genome-scale protein function classification. Bioinformatics 30:1236–1240. 10.1093/bioinformatics/btu031

Jones DT, Taylor WR, Thornton JM (1992) The rapid generation of mutation data matrices from protein sequences. Comput Appl Biosci 8: 275–282.

Kawamoto T, Noshiro M, Shen M, Nakamasu K, Hashimoto K, Kawashima-Ohya Y, Gotoh O, Kato Y (1998) Structural and phylogenetic analyses of RGD-CAP/beta ig-h3, a fasciclin-like adhesion protein expressed in chick chondrocytes. Biochim Biophys Acta 1395:288–292. 10.1016/s0167-4781(97)00172-3

Keegstra K, Talmadge KW, Bauer WD, Albersheim P (1973) The structure of plant cell walls: III. A model of the walls of suspension-cultured sycamore cells based on the interconnections of the macromolecular components. Plant Physiol 51:188–97. 10.1104/pp.51.1.188

Kieliszewski MJ (2001) The latest hype on Hyp-O-glycosylation codes. Phytochemistry 57:319–23. 10.1016/s0031-9422(01)00029-2

Kieliszewski MJ, Shpak E (2001) Synthetic genes for the elucidation of glycosylation codes for arabinogalactan-proteins and other hydroxyproline-rich glycoproteins. Cell Mol Life Sci 58:1386–1398. 10.1007/PL00000783

Kim JE, Jeong HW, Nam JO, Lee BH, Choi JY, Park RW, Park JY, Kim IS (2002) Identification of motifs in the fasciclin domains of the transforming growth factor-beta-induced matrix protein betaig-h3 that interact with the alphavbeta5 integrin. J Biol Chem 277:46159–46165. 10.1074/jbc.M207055200

Kim JE, Kim SJ, Lee BH, Park RW, Kim KS, Kim IS (2000) Identification of motifs for cell adhesion within the repeated domains of transforming growth factor-beta-induced gene, betaig-h3. J Biol Chem 275:30907–30915. 10.1074/jbc.M002752200

Knox JP, Linstead J, Peart J, Cooper C, Roberts K (1991) Developmentally regulated epitopes of cell surface arabinogalactan proteins and their relation to root tissue pattern formation. Plant J 1:317–326.

Lafarguette F, Leplé J-C, Déjardin A, Laurans F, Costa G, Lesage-Descauses M-C, Pilate G (2004) Poplar genes encoding fasciclin-like arabinogalactan proteins are highly expressed in tension wood. New Phytol 164:107–121. 10.1111/j.1469-8137.2004.01175.x

Lamport DTA (1970) Cell wall metabolism. Annu Rev Plant Physiol 21:235–70.

Ma Y, MacMillan CP, de Vries L, Mansfield SD, Hao P, Ratcliffe J, Bacic A, Johnson KL (2022) FLA11 and FLA12 glycoproteins fine-tune stem secondary wall properties in response to mechanical stresses. New Phytol 233:1750–1767. 10.1111/nph.17898

MacMillan CP, Mansfield SD, Stachurski ZH, Evans R, Southerton SG (2010) Fasciclin-like arabinogalactan proteins: specialization for stem biomechanics and cell wall architecture in Arabidopsis and Eucalyptus. Plant J 62:689–703. 10.1111/j.1365-313X.2010.04181.x

Moulia B, Coutand C, Lenne C (2006) Posture control and skeletal mechanical acclimation in terrestrial plants: implications for mechanical modeling of plant architecture. Am J Bot 93:1477–1489. 10.3732/ajb.93.10.1477

Persson S, Wei H, Milne J, Page GP, Somerville CR (2005) Identification of genes required for cellulose synthesis by regression analysis of public microarray data sets. Proc Natl Acad Sci USA 102:8633–86388. 10.1073/pnas.0503392102

Richardson KC, Jarett L, Finke EH (1960) Embedding in epoxy resins for ultrathin sectioning in electron microscopy. Stain Technol 35:313–323.

Seifert GJ (2018) Fascinating Fasciclins: A surprisingly widespread family of proteins that mediate interactions between the cell exterior and the cell surface. Int J Mol Sci 19:1628. 10.3390/ijms19061628

Shi H, Kim Y, Guo Y, Stevenson B, Zhu JK (2003) The Arabidopsis SOS5 locus encodes a putative cell surface adhesion protein and is required for normal cell expansion. Plant Cell 15:19–32. 10.1105/tpc.007872

Showalter AM (2001) Arabinogalactan-proteins: structure, expression and function. Cell Mol Life Sci 58:1399–417. 10.1007/PL00000784

Showalter AM, Keppler BD, Liu X, Lichtenberg J, Welch LR (2016) Bioinformatic identification and analysis of hydroxyproline-rich glycoproteins in Populus trichocarpa. BMC Plant Biol 16:229. 10.1186/s12870-016-0912-3

Sinnott EW (1952) Reaction wood and the regulation of tree form. Amer J Bot, 39:69–78. 10.2307/2438096

Tamura K, Stecher G, Kumar S (2021) MEGA 11: Molecular evolutionary genetics analysis version 11. Mol Biol Evol 38:3022–3027. 10.1093/molbev/msab120

Tan L, Eberhard S, Pattathil S, Warder C, Glushka J, Yuan C, Hao Z, Zhu X, Avci U, Miller JS, Baldwin D, Pham C, Orlando R, Darvill A, Hahn MG, Kieliszewski MJ, Mohnen D (2013) An Arabidopsis cell wall proteoglycan consists of pectin and arabinoxylan covalently linked to an arabinogalactan protein. Plant Cell 25:270–87. 10.1105/tpc.112.107334

Tan L, Qiu F, Lamport DT, Kieliszewski MJ (2004) Structure of a hydroxyproline (Hyp)-arabinogalactan polysaccharide from repetitive Ala-Hyp expressed in transgenic Nicotiana tabacum. J Biol Chem 279:13156–13165. 10.1074/jbc.M311864200

Thompson JD, Higgins DG, Gibson TJ (1994) CLUSTAL W: improving the sensitivity of progressive multiple sequence alignment through sequence weighting, position-specific gap penalties and weight matrix choice. Nucleic Acids Res 22:4673–4680. 10.1093/nar/22.22.4673

Wang H, Jiang C, Wang C, Yang Y, Yang L, Gao X, Zhang H (2015) Antisense expression of the fasciclin-like arabinogalactan protein FLA6 gene in Populus inhibits expression of its homologous genes and alters stem biomechanics and cell wall composition in transgenic trees, J Exp Bot 66:1291–1302. 10.1093/jxb/eru479

Wang H, Jin Y, Wang C, Li B, Jiang C, Sun Z, Zhang Z, Kong F, Zhang H (2017) Fasciclin-like arabinogalactan proteins, PtFLAs, play important roles in GA-mediated tension wood formation in Populus. Sci Rep 7:6182. 10.1038/s41598-017-06473-9

Zang L, Zheng T, Chu Y, Ding C, Zhang W, Huang Q, Su X (2015) Genome-wide analysis of the fasciclin-like arabinogalactan protein gene family reveals differential expression patterns, localization, and salt stress response in Populus. Front Plant Sci 6:1140. 10.3389/fpls.2015.01140

Zhou R, Jenkins JW, Zeng Y, Shu S, Jang H, Harding SA, Williams M, Plott C, Barry KW, Koriabine M, Amirebrahimi M, Talag J, Rajasekar S, Grimwood J, Schmitz RJ, Dawe RK, Schmutz J, Tsai CJ (2023) Haplotype-resolved genome assembly of Populus tremula × P. alba reveals aspen-specific megabase satellite DNA. Plant J 116:1003–1017. 10.1111/tpj.16454

