## Supplemental Material for "Immunolocalization of fasciclin-like arabinogalactan proteins in the G-layers of poplar tension wood fibers"

### Planta

Myiuki Takeuchi, Fernanda Trilstz Perassolo Guedes, Françoise Laurans, Nathalie Boizot, Annabelle Déjardin, Gilles Pilate.

PopFLA1
556F 5’- CACCCTGGTGCCGCGCGGCAGCATGAAGCCACAGTACTTAC -3’ (with thrombine cleavage site)
556R 5’- TTA CAG AGA AAA CAT TGC -3’

PopFLA8
560F 5’- CACCCTGGTGCCGCGCGGCAGCATGAGGCCACAATCTGTC -3’ (with thrombine cleavage site)
560R 5’- TCA TGA ATA TTT TAT TGC -3’

### Sup Table 1

**Sup Table 1** Oligonucleotide sequences used to construct the expression vectors for recombinant protein production.

| | P. tremula x P. alba_HAP1 v5.1 gene model | P. tremula x P. alba_HAP2 v5.1 gene model | | Immunogen | |
|---|---|---|---|---|---|
| | | | PopFLA8 | PopFLA1 | PF1 |
| **PopFLA1** | **PtXaTreH.19G106400** | PtXaAlbH.19G100600 | 49 | 100 | 100 |
| **PopFLA2** | PtXaTreH.13G119100 | **PtXaAlbH.13G120200** | 52 | 88 | 100 |
| **PopFLA3** | PtXaTreH.13G119200 | **PtXaAlbH.13G120300** | 49 | 84 | 100 |
| **PopFLA4** | **PtXaTreH.13G013000** | **PtXaAlbH.13G013000** | 51 | 88 | 100 |
| **PopFLA5** | **PtXaTreH.19G106500** | PtXaAlbH.19G100700 | 50 | 93 | 100 |
| **PopFLA6** | **PtXaTreH.13G119000** | PtXaAlbH.13G120100 | 49 | 83 | 100 |
| **PopFLA7** | **PtXaTreH.12G014200** | PtXaAlbH.12G015000 | 49 | 54 | 53 |
| **PopFLA8** | **PtXaTreH.09G008100** | PtXaAlbH.09G008100 | 100 | 49 | 73 |
| **PopFLA9** | PtXaTreH.04G165400 | **PtXaAlbH.04G163300** | 80 | 55 | 73 |
| **PopFLA10** | PtXaTreH.09G008000 | **PtXaAlbH.09G008000** | 46 | 44 | 47 |
| **PopFLA11** | **PtXaTreH.16G070700** | **PtXaAlbH.16G073700** | 41 | 50 | 47 |
| **PopFLA12** | **PtXaTreH.14G131800** | PtXaAlbH.14G133100 | 32 | 33 | 27 |
| **PopFLA13** | PtXaAlbH.06G112300 | **PtXaAlbH.06G112300** | 43 | 50 | 47 |
| **PopFLA14** | PtXaAlbH.01G264300 | **PtXaAlbH.01G264300** | 48 | 52 | 53 |
| **PopFLA15** | **PtXaTreH.15G101700** | PtXaAlbH.15G102200 | 58 | 58 | 53 |

### Sup Table 2

**Sup Table 2** Identity at the amino acid level between the fifteen PopFLAs expressed in the xylem and the three sequences used as immunogens to produce each of the 3 antibodies directed against the PopFLA1, PopFLA8 and PF1 peptides. For each PopFLA, the gene models for both the *Populus tremula* and the *Populus alba* alleles are indicated according to the genome sequences available at https://phytozome-next.jgi.doe.gov/. In bold, the haplotype(s) that are 100% homologous to the transcripts present in xylem tissues; red: sequences with 73 to 100% identity between the PopFLA sequence and the immunogen sequence; green: 41 to 58% identity.

**Sup Fig. 1** Phylogenetic tree built via the tree joining method in MEGA. The tree with the highest log likelihood (-10872,55) is shown. The data are drawn to scale, with branch lengths measured in terms of the number of substitutions per site. This analysis involved 45 amino acid sequences: AtFLA11 and AtFLA12 sequences, 15 PopFLAs identified in *P. tremula × P. alba* wood samples (Lafarguette et al., 2004) and the 28 PtFLAs from *P. trichocarpa* with the highest homology with AtFLA11 and AtFLA12 (Showalter et al., 2016). There was a total of 301 positions in the final data set. Blue frame: PopFLA1 and PopFLA8 used as immunogens for antibody production; red frame: the 13 remaining PopFLAs; green frame AtFLA11 and AtFLA12.

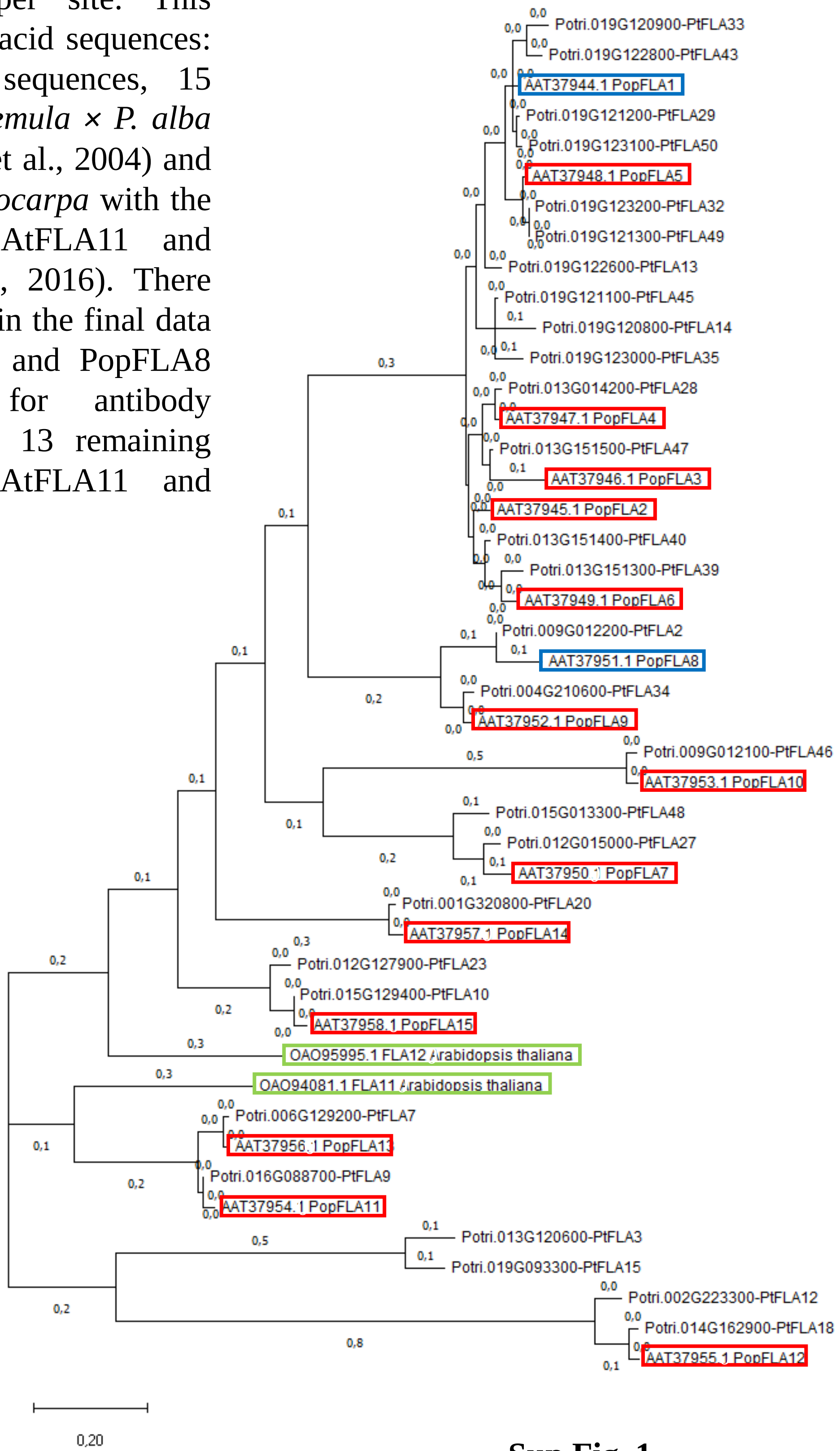


**Sup Fig. 1**

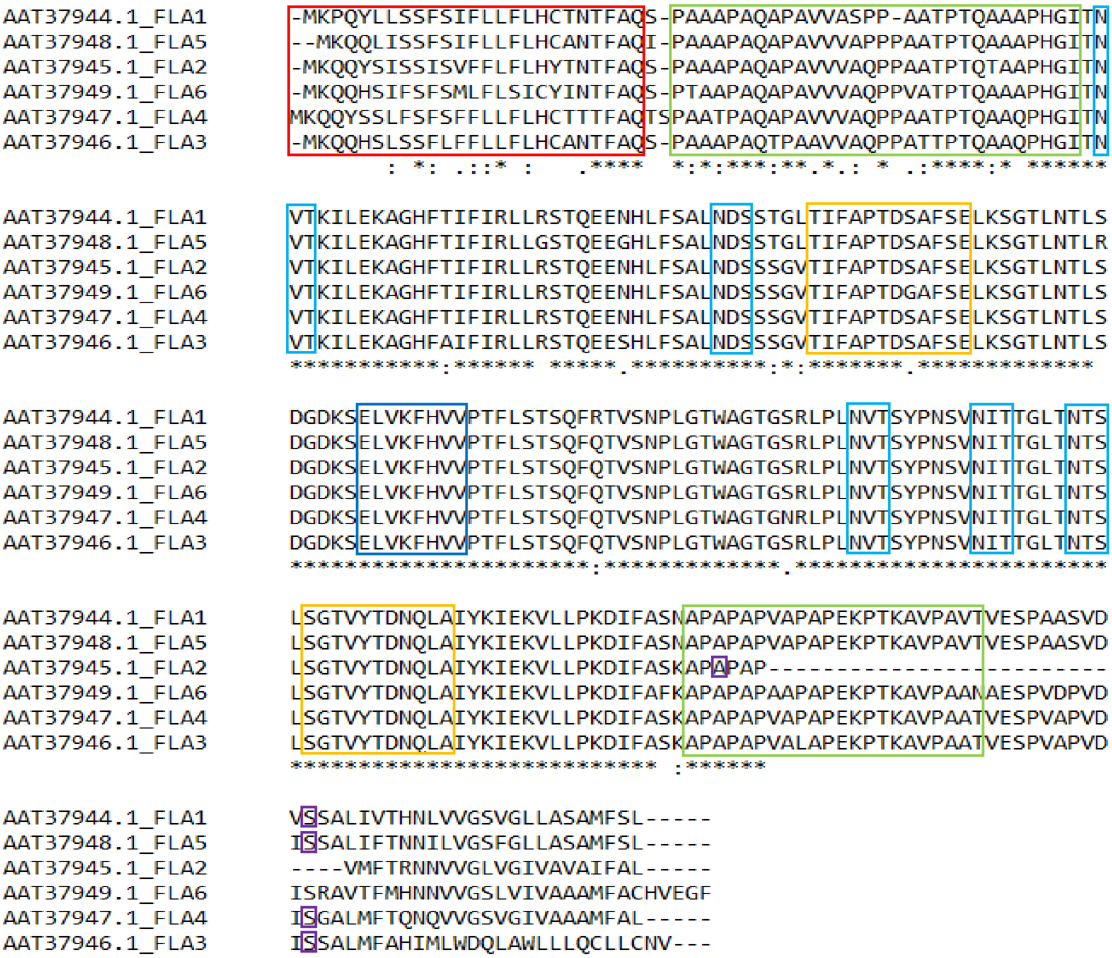


**Sup Fig. 2**

**Sup Fig. 2** Sequence alignment of PopFLA1 to PopFLA6. Red frame: signal peptide for secretion; green frames: AGP-like domains; yellow frames: HI and H2 conserved regions characteristic of the FAS1 domain; dark blue frame: amino acids involved in adhesion; light blue frames: N-glycosylation sites; purple frame: positions of GPI anchor signal sequences not present in PopFLA6.

```
a   PopFLA11      DQQKVQLVQFHILPN
    PopFLA13      DQQKVQLVQFHIIPN
    PopFLA1       DGDKSELVKFHVVPT
    PopFLA2       DGDKSELVKFHVVPT
    PopFLA3       DGDKSELVKFHVVPT
    PopFLA4       DGDKSELVKFHVVPT
    PopFLA5       DGDKSELVKFHVVPT
    PopFLA6       DGDKSELVKFHVVPT
    PopFLA8       DEDKTELVKFHVLPA
    PopFLA9       DEDKTELVKFHVLPA
    PopFLA14      DQEKVELMQFHIVPM
    PopFLA15      DQEKAELVQFHIIPQ
    PopFLA10      DREKLEFVQFHILPR
    PopFLA7       DHQKIELVQFHIIPK
    PopFLA12      QDQLKQLILFHALPH
                  : :  ::: ** :*

b   PopFLA1       DGDKSELVKFHVVPT
    PopFLA2       DGDKSELVKFHVVPT
    PopFLA3       DGDKSELVKFHVVPT
    PopFLA4       DGDKSELVKFHVVPT
    PopFLA5       DGDKSELVKFHVVPT
    PopFLA6       DGDKSELVKFHVVPT
    PopFLA8       DEDKTELVKFHVLPA
    PopFLA9       DEDKTELVKFHVLPA
                  * **:*******:*:
```

**Sup Fig. 3**

**Sup Fig. 3** Multiple alignment of the PF peptide (15 aa) among the 15 PopFLAs found in poplar xylem (a) and the 8 most homologous TWs specific to PopFLA (b). * conserved amino acid among all the aligned sequences;  : conservative substitution
